# Comparative proteome analysis revealed the differences in responses of bovine alveolar macrophages to both M.tb and Mb infection

**DOI:** 10.1101/2021.06.16.448489

**Authors:** Yurong Cai, Yong Li, Weifeng Gao, Pu Wang, Gang Zhang, Lingling Jiang

## Abstract

Bovine tuberculosis, caused by both MTB(Mycobacterium tuberculosis) and MB(Mycobacterium bovis), is one of the most serious zoonotic diseases in the world. Both pathogenic MTB and MB interact closely with the host alveolar macrophages. However, the mechanisms of defense response of macrophages to MTB or MB still need further researches. Presently, in-depth comparative proteome analysis was performed on the MB- and MTB- infected macrophage. We identified that macrophages was more sensitive to the MB infection. Both pathogens could induce strong energy metabolism of macrophages. To eliminate MB and protect the host, a bunch of proteins and signaling which involved in autophagy- and inflammatory-related progresses were highly activated in macrophages following MB infection. In contrast, only proteins relevant to energy metabolisms were highly expressed in macrophages following MTB infection. In summary, we proposed that macrophages were more sensitive to MB attacks and could deployed proteins functioned in autophagy- and inflammatory-related progresses to protect the host. Our research provide a novel insight into the different mechanisms of defense responses of macrophages to both MTB and MB infection.

## Introduction

Tuberculosis (TB) is a notorious infectious disease that has been documented for a profound time. It was once the disease which caused a large amount of death of homo sapiens and various animals, and was also known as the “White Plague.” Currently, tuberculosis is still one of the most serious infectious diseases in the world and with the increase of population mobility (Li ,2011). Such serious disease is chronic infectious disease caused by the infection of Mycobacterium tuberculosis and Mycobacterium bovis, which can attack both cattle and Homo sapiens (Daley et al., 2010). Therefore, tuberculosis not only hinders the development of cattle breeding industry, but also seriously endangers human life and health. An overall survey showed that the global number of new tuberculosis cases in 2016 was 10.4 million, among which about 10% of human Tuberculosis cases were caused by the infection of Mycobacterium bois infection (Waters and PALMER 2015; Minae et al., 2017). Hence, effective control of bovine Tuberculosis will be a new strategy for reducing the risk of human infection with Tuberculosis. However, the mechanisms of bovine resistance to the infection of Mycobacterium tuberculosis and Mycobacterium bovis are still not clear.

Macrophages, as immune regulatory cells and effector cells, regulate the inflammatory response and immune response of the body through phagocytosis, antigen presentation and secretion of various cytokines during the process of infection. Such cells are imperative immune cells of the body and play important roles in host anti-infection responses (Jing et al., 2016; Hmama et al., 2015). Mycobacterium bovis, as a typical intracellular parasite, macrophages can provide MTB with nutrients and places for survival and reproduction, and can also eliminate MTB through apoptosis or autophagy (Moraco and Kornfeld, 2014). Well studied is that MTB can prevent phagocytic acidification, phagocytic and lysosomal fusion, so as to avoid proteolytic enzyme hydrolysis and subsequent immune response events, which is the main strategy for MTB to avoid host cell clearance (Horacio et al., 2008; Wel et al., 2007). As the ability to infect macrophages is crucial for the transmission and diffusion of pathogenic bacteria in the host, the interaction between MB and macrophages has been the focus in the study of anti-tuberculosis immune mechanism. Various studies (Fu et al., 2017; Yi et al., 2014; Yang et al., 2016) have shown that MB infection can cause abnormal expression of long non-coding RNA and mRNA expression profile in the body, and these differentially expressed lncRNAs are involved in the regulation of cell signaling pathways such as Toll like receptor signaling pathway (TLR), Transforming Growth factor-βsignaling pathway (TGF-β) and Hippo signaling pathway (HPO). During the infection of Mycobacterium tuberculosis, there is specificity in the immune response of organism to pathogen (Masters et al., 2010). Cattle themselves serve as hosts for Mycobacterium tuberculosis infection and are ideal large animal models for tuberculosis research. Therefore, researches on the interaction between bovine macrophages and Mycobacterium tuberculosis will contribute to clarify the mechanism of host resistance to tuberculosis infection.

In the process of immune response, Macrophages plays a crucial role to fight against pathogenic bacteria infection, control the development of the tissue inflammation (Wynn etal 2013). Alveolar macrophages are an important barrier to defense the infection of with various methods, it has been clearly proved that Macrophages could defense MB infection by phagosome fusion with lysosomes, antigen presented to the start of the immune response, activation of toll-like receptor, Macrophage cells apoptosis and destruction the secretion of cytokines. Macrophages will degrade the phagocytic MTB through acidic hydrolase in lysosomes, thus killing or inhibiting its endogenous growth (Wang et al. 2010). Meanwhile, as an important antigen transmitting cell, macrophages can degrade antigens into immunogenic peptides through endocytosis, and then produce and release IFN-Y to inhibit the growth of MB in the cell (Canaday et al., 2001). Macrophages can also enhance the toxicity of reactive oxygen products and reactive nitrogen products and stimulate the formation of phagocytic lysosomes to further defense the diffusion of MB (Sureka et al., 2010). Additionally, the activation of TNF-α, IL-1, IL-6, IL-15, IL-10, IL-12 and NF-kβ also plays important roles in the defense response of Macrophages to MB infection (Sánchez et al., 2012).

In the present study, we identified the differences in responses of Macrophages to MTB and MB infection, through comparison proteomic analysis, and deciphered the mechanisms of MB-Macrophage interaction. We found that autophagy- and inflammatory-related progresses were significantly influenced by both MB- and MTB-infection in macrophage cells. In parallel, various signaling pathways relevant to auto autophagy and inflammatory were altered by MB-infection, whereas MTB-infection did not significantly invoked their activation. The output of this our research, regarding the complex specific response of macrophage following MB challenges, and contribute out understanding of the interaction progresses between MB-Macrophage in establishing the host immune response to MTB.

## Materials and Methods

### Bacterial Strains and Culture Conditions

The M.tb clinical strain Gong( M.tb1 ), Wu( M.tb2 ), Zhang( M.tb3 ), and M. bovis 1054( Mb1 ) 1060( Mb2 )1087( Mb3 )were kindly provided by Dr. Xiaopin Wang, The fourth people’s hospital of Ningxia Hui Autonomous Region(Yinchuan, China). All strains were cultured to mid-log phase in Middlebrook 7H9 medium (Becton Dickinson and Company) supplemented with 10% catalase medium (OADC) (Becton Dickinson and Company) and 0.2% Tween 80 (Biotopped life sciences). Cultures were maintained in a biosafety level 3 facility at The fourth people’s hospital of Ningxia Hui Autonomous Region(Yinchuan, China) and stored at −80°C. Bacteria were harvested from the culture medium by centrifugation at 3000 rpm for 10 min washed once in phosphate-buffered saline (PBS) (Biological Industries , BI)), resuspended in PBS to an OD600 of 1, equivalent to 3 × 108 bacteria/ml.

For the colony counting assay, bacteria were serially diluted 10-fold with Middlebrook’s 7H9 medium and 100μL of each dilution was transferred to Middlebrook 7H10(Becton Dickinson and Company) plates supplemented with 10% catalase medium (OADC) (Becton Dickinson and Company), 0.5% oleic acid( Solarbio life science ), and 0.05% Tween 80 (Biotopped life sciences). Bacteria were grown for 8-10 weeks until the colonies bacterial lawns grow. Bacteria were harvested from the bacterial lawns, resuspended in PBS to an OD600 of 1 obtained to guarantee the same initial infection dose.

### Cell Collection, Culture and Infection

Primary bovine alveolar macrophages were collected from a beef cattle slaughterhouse in Ningxia(Yinchuan, China). The fresh healthy bovine lungs were infused into the lungs via trachea with physiological saline supplemented with 4% antibiotic-antimycotic(100×)( SolarBio Life Science). The lavage fluid containing bovine alveolar macrophages(BAM) was collected, passed through a nylon mesh(75μm), centrifugation at 10,000 g for 5 min at 25℃. The pellet is lysed with Red Blood Cell Lysis Buffer (solarbio life csience) for 10 minutes at room temperature, Centrifuge again. Rinse 3 times with PBS, resuspend the pellet in DMEM supplemented with 10% FBS and 4% antibiotic-antimycotic and incubated at 37°C in a 5% CO2 atmosphere. To reduce the effect of antibiotic-antimycotic from BAM, the medium was replaced, and incubation was continued for 6h,12h, and 24h. The medium supplemented with 0.5% antibiotic-antimycotic at 24h.

The medium supplemented with 10% FBS was replaced, at 36h.They were then infected with M.tb clinical strain Gong(M.tb1), Wu(M.tb2), Zhang(M.tb3), and M. bovis 1054(Mb1), 1060(Mb2), 1087(Mb3)at an MOI of 10(10 bacteria to one cell). After 6 h, cells were collected, washed one time with ice-cold PBS and processed for whole protein extraction. Three independent experiments were performed.

### Sample preparation for label-free proteomic quantification

Sample was sonicated three times on ice using ultrasonic processor in lysis buffer containing 8 M urea, 1% Protease Inhibitor Cocktail. The remaining debris was removed by centrifugation at 12,000 g at 4□ for 10 min. Finally, the supernatant was collected and the protein concentration was determined with BCA kit according to the manufacturer’s instructions. For digestion, the protein solution was reduced with 5 mM dithiothreitol for 30 min at 56□ and alkylated with 11 mM iodoacetamide for 15 min at room temperature in darkness. The protein sample was then diluted by adding 100 mM TEAB to urea concentration less than 2M. Finally, trypsin was added at 1:50 trypsin-to-protein mass ratio for the first digestion overnight and 1:100 trypsin-to-protein mass ratio for a second 4 h-digestion. Lastly, the resulting peptides were washed, eluted, dried and re-solubilised in 20 μl of buffer containing 3% acetonitrile and 0.1% formic acid for subsequent Liquid Chromotrophy Mass Spectrometry analysis.

### Protein Identification by LC-MS/MS

The tryptic peptides were dissolved in 0.1% formic acid, directly loaded onto a home-made reversed-phase analytical column. The gradient was comprised of an increase from 6% to 23% solvent 0.1% formic acid in 98% acetonitrile over 26 min, 23% to 35% in 8 min and climbing to 80% in 3 min then holding at 80% for the last 3 min, all at a constant flow rate of 400 nL/min on an EASY-nLC 1000 Ultra Performance Liquid Chromatography system. The peptides were subjected to NSI source followed by tandem mass spectrometry (MS/MS) in Q ExactiveTM Plus (Thermo) coupled online to the UPLC. The electrospray voltage applied was 2.0 kV. The m/z scan range was 350 to 1800 for full scan, and intact peptides were detected in the Orbitrap at a resolution of 70,000. Peptides were selected using NCE setting as 28 and the fragments were detected in the Orbitrap at a resolution of 17,500. A data-dependent procedure that alternated between one MS scan followed by 20 MS/MS scans with 15.0s dynamic exclusion.

### Data analysis

The resulting MS/MS data were processed using Maxquant search engine (vs 1.6.3.3, Cox and Mann, 2008). Tandem mass spectra were searched against the uniprot database concatenated with reverse decoy database. Trypsin/P was specified as cleavage enzyme allowing up to 4 missing cleavages. The mass tolerance for precursor ions was set as 20 ppm in First search and 5 ppm in Main search, and the mass tolerance for fragment ions was set as 0.02 Da. Carbamidomethyl on Cys was specified as fixed modification and acetylation modification and oxidation on Met were specified as variable modifications. P-value was adjusted to < 5% and minimum score for modified peptides was set > 40.

### Subcellular localization prediction

we used wolfpsort a subcellular localization predication soft to predict subcellular localization. Special for protokaryon species, Subcellular localization prediction soft CELLO was used.

### Enrichment analysis of Gene Ontology

Proteins were classified by GO annotation into three categories: biological process, cellular compartment and molecular function. For each category, a two-tailed Fisher’s exact test was employed to test the enrichment of the differentially expressed protein against all identified proteins. The GO with a corrected p-value < 0.05 is considered significant.

### Pathway enrichment analysis

Encyclopedia of Genes and Genomes (KEGG) database was used to identify enriched pathways by a two-tailed Fisher’s exact test to test the enrichment of the differentially expressed protein against all identified proteins. The pathway with a corrected p-value < 0.05 was considered significant. These pathways were classified into hierarchical categories according to the KEGG website.

### Protein-protein Interaction Network

All differentially expressed protein database accession or sequence were searched against the STRING database version 10.1 for protein-protein interactions. Only interactions between the proteins belonging to the searched data set were selected, thereby excluding external candidates. STRING defines a metric called “confidence score” to define interaction confidence; we fetched all interactions that had a confidence score ≥ 0.7 (high confidence). Interaction network form STRING was visualized in Cytoscape 3.0.

qRT-PCR verification of gene expression of autophagy-related proteins

The expression of important defense- and autophagy-related proteins induced by MB infection (Q0VCQ6, Q05204 and Q8HXK9) was investigated through real-time quantitative polymerase chain reactions (RT-qPCR). The specific primers for RT-qPCR were designed using Primer 6(v6.24) Designer.

Total RNA extraction and quantification were performed as described in Vieira et al.(2016) using the Rneasy MiNi Kit (Qiagen, Germany) and the NanoDrop 2000 Spectrophotometer (Thermo Scientific, Waltham, MA, United States), respectively. cDNA was synthesized with a RevertAidTM First Strand cDNA Synthesis Kit following the manufacturer’s protocol (Qiagen, Germany). All the reactions were carried out three times for three independent biological replicates. The relative transcript levels of the target genes was calculated using the foldchange of 2^-△△Ct^ value(Schmittgen et al.2008). Primer sequences for qRT-PCR are shown in Table 1.

**Table 1.**
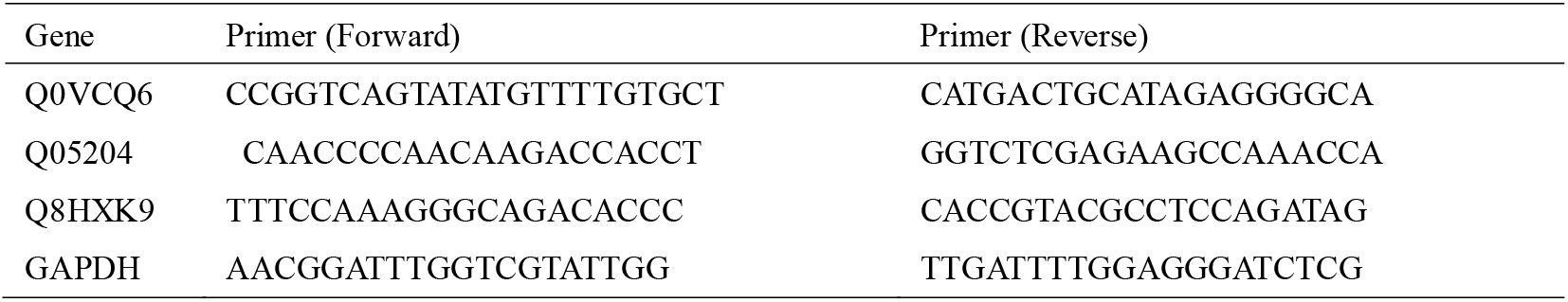
Primer sequence for real-time PCR

## Results

### Quantitative Proteomic Analysis by Label Free

To investigate the changes in proteomic profiles of BAM, which was caused by the infection of MTB and MB, total proteins of normal BAM (control), MB- and MTB-infected BAM with three replicates were extracted, and then analyzed by LC/ESI-MS/MS and quantified by Label Free. Totally, 46 018 spectra were generated, and 5467 proteins were identified against the bovine reference.

The two score plots of the PCA models show a clear separation of samples from different experimental groups (MTB and MB) and controls groups BAM, accounting for 36.6% of the observed variance, indicating that pathogenic bacteria infection was the key factor affecting protein expression (Figure 1A). A genotype effect was also observed explaining about 21.6% of variance (Figure 1A). Remarkably, the MB-infected samples showed a slight interaction with other two sample groups (MTB and BAM), suggesting that under MB infection challenge BAM could make some similar responses with MTB challenges (Figure 1A). Additionally, correlation analysis displayed a well consensus with the PCA analysis that the variations of the three biological replicates were calculated according to their quantitative data, and they all showed little variation between three biological replicates, indicating a better quality and reproducibility of the data (Figure 1B). Therefore, we proposed that MB-infection and MTB-infection will all result in significant changes in gene expression pattern of BAM, and some similar response could be invoked by both MB-infection and MTB infection. The same and different responses caused by both pathogenic bacteria will be the focus below.

**Figure 1.**
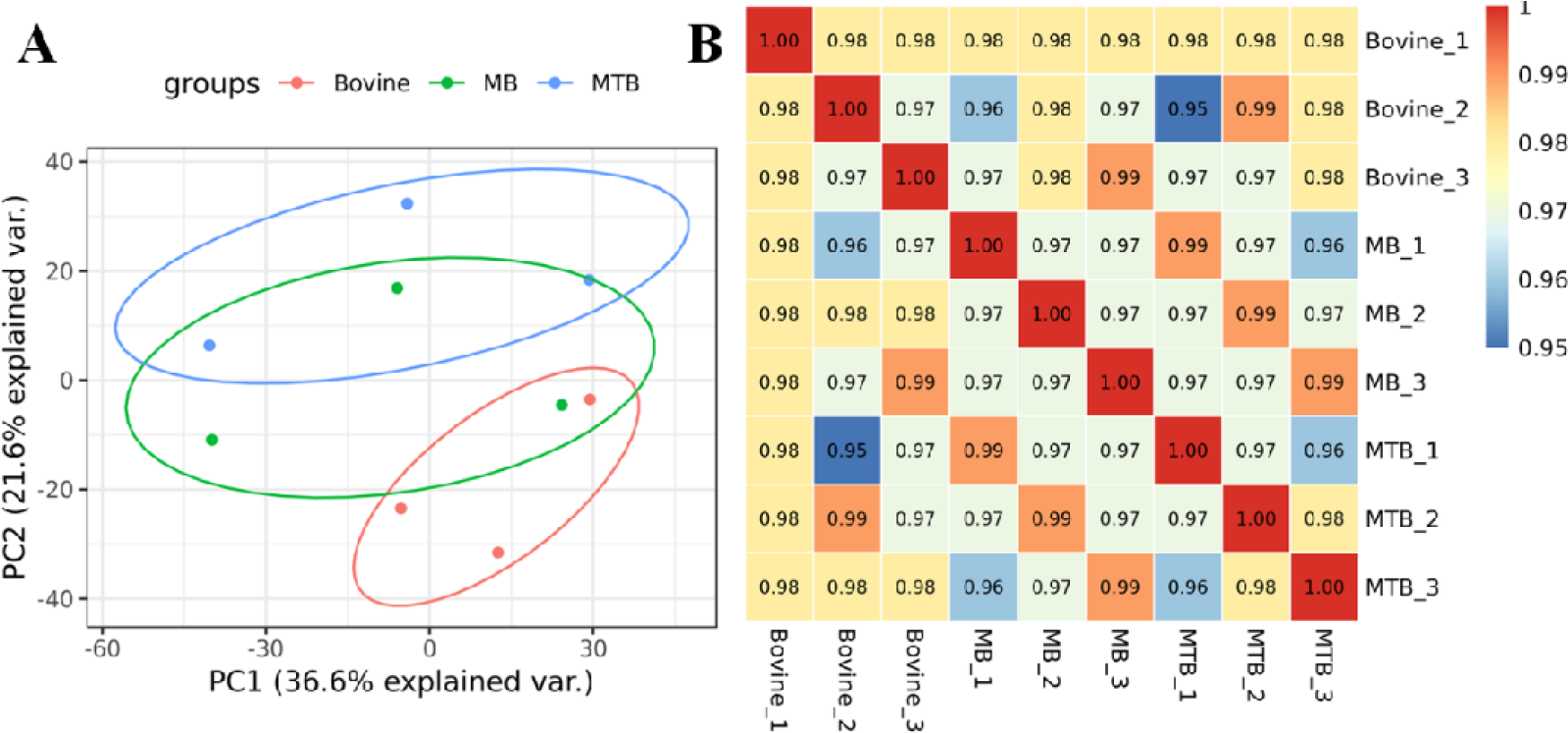
Evaluation of integral proteomic profiles from three treatment. A: Principal component (PC) analysis of protein profiling data from MB-infected, MTB-infected and control BAM. Each replicate with the same treatment was shown in the same plots (red: Bovine; green: MB; blue: MTB). PC1 for proteins explains 36.6% of variance and PC2 explains 21.6% of variance using integral proteomic data. B: Correlation analysis on integral protein data from three treatment. The pearson value of correlation was shown in the cells.

### Differentially expressed proteins in the bovine alveolar macrophages after being challenged by the MTB- and MB-infection

To determine the in-depth variation between MTB-infection and MB-infection, two pairwise comparisons were designed, including MTB.vs.BAM and MB.vs.BAM. In total, 18 proteins were significantly up or down-regulated during MTB infection(Figure 2A, B; Log1.5FC < −1 or >1, P < 0.05). We identified that 17 proteins were up-regulated and 1 protein was down-regulated during MTB infection (Figure 2A). Only one host protein, regulator Prefoldin subunit 5 (Q8HYI9), was down-regulated by MTB infection. In parallel, 60 proteins were up-regulated and 3 proteins were down-regulated during MB infection (Figure 2B). The three down-regulated proteins were TBC1 domain family member 2A, Polysaccharide biosynthesis domain containing 1 and Beta-defensin 10 (A6QP29, F1MV85 and P46168; Table 5).

**Figure 2.**
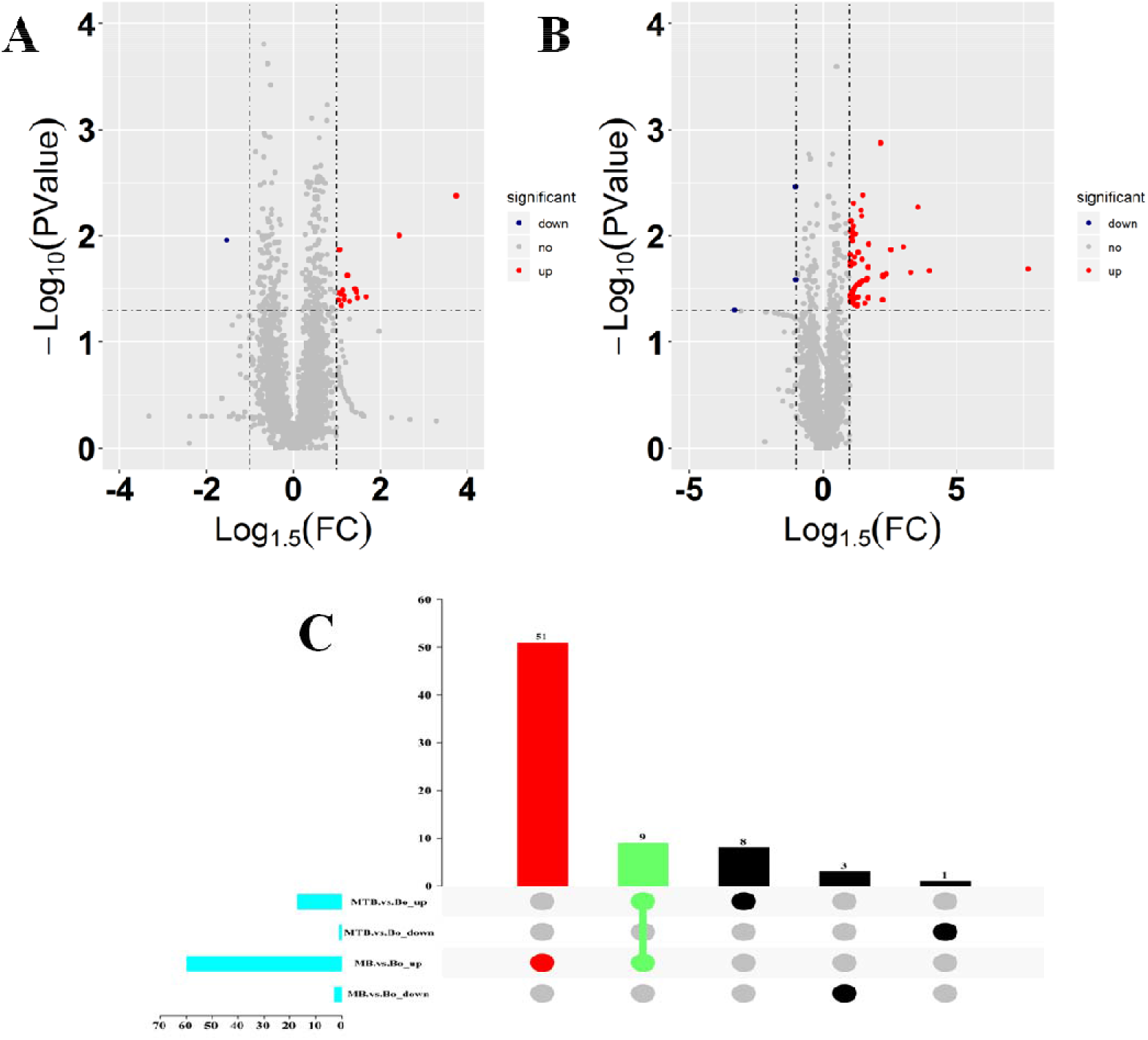
Identification of differentially expressed proteins in both MB.vs.BAM and MTB.vs.BAM pairwise comparisons. A, B: Volcano plots for the potential MTB-(A) and MB-induced (B) metabolomic features of BAM. Red points indicate significantly up-regulated proteins between the two groups (Log1.5FC < −1.0 or > 1.0; q-value < 0.05). The blue points showed the significantly down-regulated proteins between the two groups. The gray points showed tentatively matched features with no significance. C: Up-set diagrams representing the overlap of identified differentially expressed proteins in both MB.vs.BAM and MTB.vs.BAM pairwise comparisons, which were up-regulated in any two groups and specifically differentially expressed in MB-and MTB-infected BAM.

In addition, to show the proteins which were up-regulated or down-regulated during MTB-and MB-infection BAM, up-set diagrams were shown as Figure 2C. Among these up-regulated proteins, we further noted that 9 up-regulated proteins were shared by MTB- and MB-infection (Figure 2C). Meanwhile, 51 proteins were only invoked to up-regulate by MB-infection and 8 proteins could be induced by MTB-infection (Figure 2C). The results indicated that MB-infection could stimulate the BAM to make more responses than MTB-infection. However, no down-regulated proteins were shared by both MTB.vs.BAM and MB.vs.BAM (Figure 2C). Based on this, three kinds of up-regulated proteins will be the analysis focus below to elucidate the similarities and differences in response of MTB- and MB-infected bovine cells.

### Functional annotation based on GO and KEGG analysis on nine MTB- and MB-induced up-regulated proteins

In order to detect whether the nine up-regulated proteins were significantly enriched in certain functional types, Gene Ontology (GO) and KEGG enrichment analyses were performed on these proteins (Figure 3; Table 2). To clearly investigate the similarities between MTB.vs.BAM and MB.vs.BAM pairwise comparisons, we firstly performed GO enrichment analysis on these nine up-regulated proteins (Figure 3A, B, C; Table 2). We found that terms relevant to biological progresses were the most significant enriched categorizes, whereas terms involved in cellular component and molecular function were relatively less (Figure 3A, B, C; Table 2). Being challenged by both MTB and MB pathogenic bacteria, BAM will activated various terms relevant to cellular component, including lysosomal lumen, lytic vacuole, lysosome and autophagosome (Figure 3A; Table 2). It has been reported that lysosome-related proteins were involved in defense response of macrophage to MTB and MB infection (Bach et al., 2008; Wel et al., 2007). We further noted that many immunity-related terms which involved in molecular function were activated in response MTB- and MB-infection, including NADPH-hemoprotein reductase activity, interleukin-1 receptor binding, scavenger receptor activity, calcium-dependent protein binding and cytokine activity (Figure 3B; Table 2). As being attacked by pathogenic bacteria, calcium-related signaling pathways will rapidly transmit the signal to activate the defense response, including a start of an inflammatory reaction (A. Casadevall et al., 2008; L. Ramakrishnan et al., 2014). Importantly, in biological progress categorizes, many defense-related terms were significantly invoked by both pathogenic bacteria infection, including acute-phase response, cellular response to molecule of bacterial origin, cellular response to biotic stimulus, response to molecule of bacterial origin, acute inflammatory response, inflammatory response, defense response, response to bacterium, cellular response to oxygen-containing compound, regulation of antimicrobial humoral response, positive regulation of antimicrobial humoral response, response to biotic stimulus and regulation of inflammatory response (Figure 3C; Table 2). Together, we concluded that the autophagy-related terms, inflammatory-related progresses and defenses to pathogenic bacteria were the important responses of BAM, which was invoked by the MTB and MB attack challenges.

**Figure 3.**
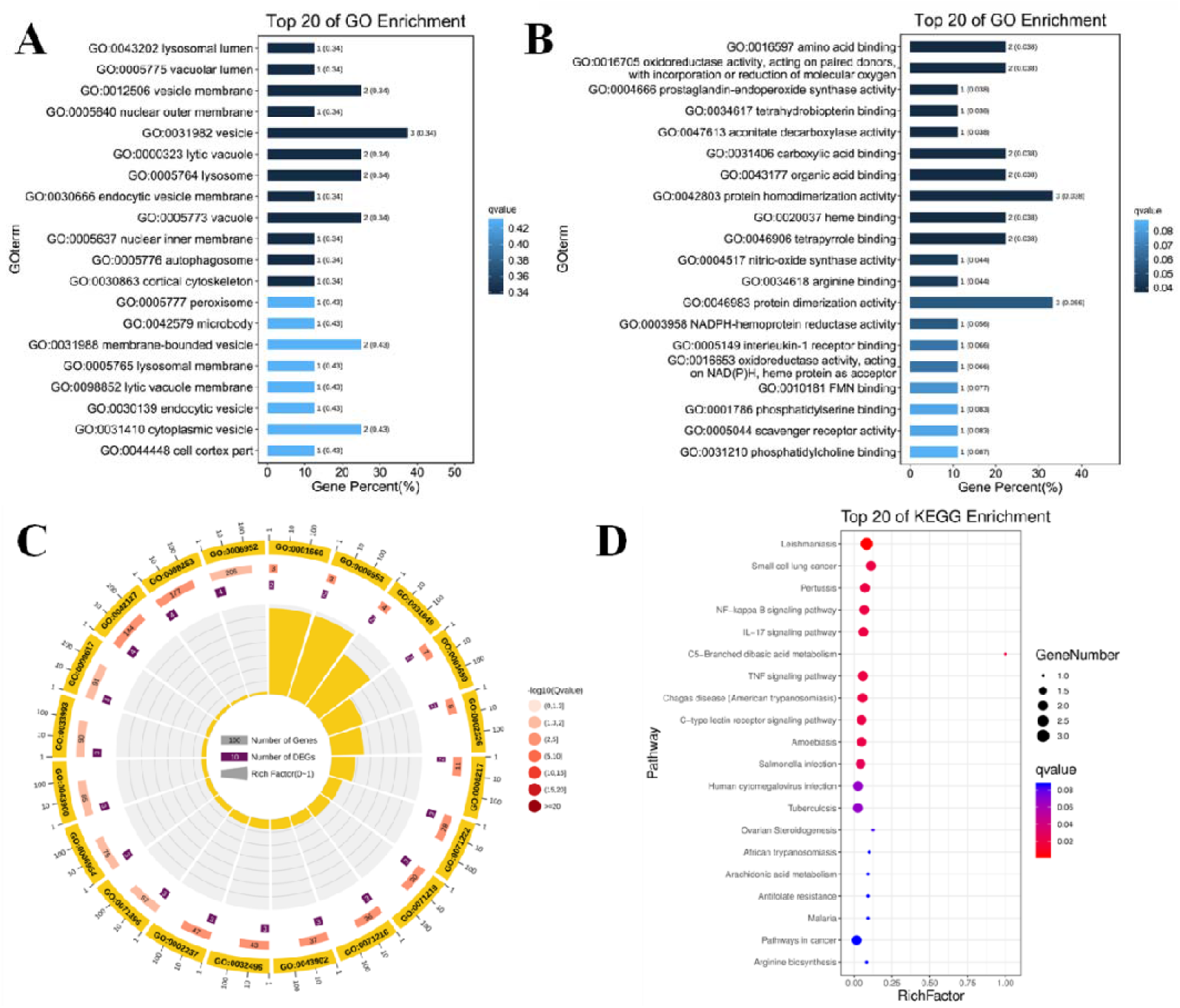
Functional annotation analysis on 9 MTB- and MB-induced up-regulated proteins. A, B, C: GO categories assigned to bovine proteins. The proteins were categorized according to the annotation of GO, and the number of each category is displayed based on biological process (C), cellular components (A), and molecular functions (B).The number of proteins involved in each GO category were displayed on each terms. The color of each column represented the significance of each enriched categorize. D: The pathway enrichment analysis for these significant 9 proteins in MB-and MTB-infected BAM. The scatter plot visualized the pathway impact and enrichment results for all these 9 significantly up-regulated proteins. Each point represented different metabolic pathways. The various color levels displayed different levels of significance of metabolic pathways from low (blue) to high (red). The different sizes of each point were used to represent the number of proteins involved in the corresponding metabolic pathway. Moreover the corresponding pathway name of each point is labeled.

**Table 2.** Significantly enriched Gene Ontology terms in both MTB-and MB-infected bovine alveolar macrophages.

| GO ID | Description | Gene Number | pvalue | Genes |
| --- | --- | --- | --- | --- |
| GO:0043202 | lysosomal lumen | 1 | 0.010493 | A0A3S5ZPN6 |
| GO:0000323 | lytic vacuole | 2 | 0.030988 | A0A3S5ZPN6; P09428 |
| GO:0005764 | lysosome | 2 | 0.030988 | A0A3S5ZPN6; P09428 |
| **GO:0005776** | autophagosome | 1 | 0.048991 | P09428 |
| **GO:0003958** | NADPH-hemoprotein<br>reductase activity | 1 | 0.009069 | F1MYR5 |
| **GO:0005149** | interleukin-1 receptor binding | 1 | 0.012076 | P09428 |
| **GO:0005044** | scavenger receptor activity | 1 | 0.018066 | A0A3S5ZPN6 |
| **GO:0048306** | calcium-dependent protein<br>binding | 1 | 0.026989 | F1MX83 |
| **GO:0005125** | cytokine activity | 1 | 0.041702 | P09428 |
| **GO:0006953** | acute-phase response | 2 | 1.04E-05 | F1MNI5; P09428 |
| **GO:0071219** | cellular response to molecule<br>of bacterial origin | 3 | 1.87E-05 | F1MGW6; F1MYR5;<br>P09428 |
| **GO:0071216** | cellular response to biotic<br>stimulus | 3 | 3.27E-05 | F1MGW6; F1MYR5;<br>P09428 |
| **GO:0002237** | response to molecule of<br>bacterial origin | 3 | 7.37E-05 | F1MGW6; F1MYR5;<br>P09428 |
| **GO:0002526** | acute inflammatory response | 2 | 9.64E-05 | F1MNI5; P09428 |
| **GO:0006954** | inflammatory response | 3 | 0.0003 | F1MNI5; F1MYR5;<br>P09428 |
| **GO:0006952** | defense response | 4 | 0.000312 | F1MGW6; F1MNI5;<br>F1MYR5; P09428 |
| **GO:0009617** | response to bacterium | 3 | 0.000534 | F1MGW6; F1MYR5;<br>P09428 |
| **GO:1901701** | cellular response to<br>oxygen-containing compound | 3 | 0.001038 | F1MGW6; F1MYR5;<br>P09428 |
| **GO:0002759** | regulation of antimicrobial<br>humoral response | 1 | 0.002037 | F1MGW6 |
| **GO:0002760** | positive regulation of<br>antimicrobial humoral<br>response | 1 | 0.002037 | F1MGW6 |
| **GO:0009607** | response to biotic stimulus | 3 | 0.002496 | F1MGW6; F1MYR5;<br>P09428 |
| **GO:0050727** | regulation of inflammatory<br>response | 2 | 0.00301 | F1MNI5; F1MYR5 |

***“P < 0.05”***

To further investigate the role of up-regulated proteins in response to MTB and MB infection, these nine significantly up-regulated proteins from MTB- and MB-infected BAM were further annotated by KEGG pathway analysis with p < 0.05 against animal reference pathways from KEGG database (Figure 3D). Up-regulated of Label Free quantitative proteins after infection of both pathogenic bacteria was mainly enriched in NF-kappa B signaling pathway, IL-17 signaling pathway, C-type lectin receptor signaling pathway, Tuberculosis, Cytokine-cytokine receptor interaction, Inflammatory mediator regulation of TRP channels, Toll-like receptor signaling pathway, Peroxisome and HIF-1 signaling pathway (Figure 3D). These pathways mainly performed functions in inflammatory- and immunity-related responses of host cells as being attacked by pathogenic microbial. It has been proved that NF-kappa B signaling pathway, IL-17 signaling pathway and Toll-like receptor signaling pathway have important function in tuberculosis (A. Casadevall et al., 2008; L. Ramakrishnan et al., 2014; Yang et al., 2016; Yi et al., 2014; Fu et al., 2017). Therefore, we highlight that both MTB and MB infection could cause inflammatory responses of BAM.

### Functional analysis on up-regulated proteins which were specifically activated by MTB infection

Gene ontology analysis provides a generally accepted identification set to describe proteins attributes in an organism. Hence, we used GO functional annotation analysis on the up-regulated proteins which were only induced to over-express by MTB to elucidate the specific response of BAM under MTB challenges (Figure 4A; Table 3). All these eight MTB-induced proteins were annotated with categories simultaneously. The classical three terms, molecular functions, biological processes and cellular components, were widely covered (Figure 4A; Table 3). The results showed that the terms involved in molecular functions and cellular components were the main categorizes which invoked to significantly enriched in BAM by MTB infection (Figure 4A; Table 3). For cellular component category, fourteen terms were significantly enriched, including fibrinogen complex, mitochondrial respiratory chain complex I, NADH dehydrogenase complex, respiratory chain complex I, mitochondrial respiratory chain, respiratory chain complex, respiratory chain, oxidoreductase complex, photoreceptor inner segment, inner mitochondrial membrane protein complex, mitochondrion, mitochondrial protein complex, mitochondrial part and mitochondrial membrane part (Figure 4A; Table 3). The results further showed that all significantly enriched cellular component-related terms were involved in energy metabolism and performed functions in mitochondria, suggesting that the MTB infection will induce dramatic changes in energy metabolisms of BAM, which could help host cells to defense pathogenic bacteria attacks. Similarly, in molecular function category, the terms relevant to energy metabolisms were also the active terms of BAM in response to MTB infection, including NADH dehydrogenase activity, oxidoreductase activity, acting on NAD(P)H, lysophosphatidic acid phosphatase activity, oxidoreductase activity, acting on NAD(P)H, inorganic diphosphatase activity, aldehyde dehydrogenase (NADP+) activity and NADP-retinol dehydrogenase activity (Figure 4A; Table 3). Therefore, we proposed that as being challenged by MTB infection, BAM will accelerate energy metabolism to defense the MTB attacks to protect the organisms.

**Figure 4.**
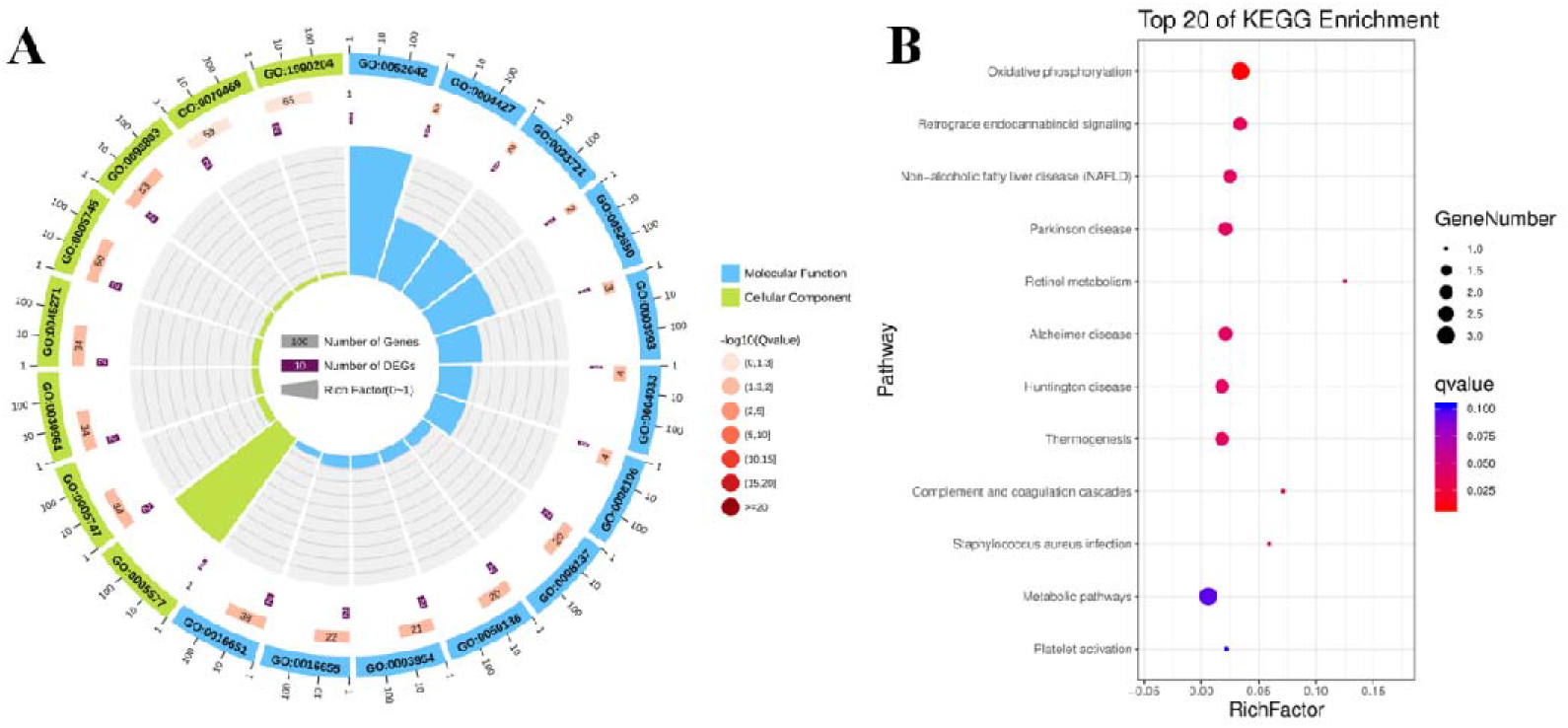
Functional annotation analysis on the up-regulated proteins which were only activated by MTB infection. A: GO categories analysis on the up-regulated proteins associated with the MTB infection. The proteins were categorized according to the annotation of GO, and the number of each category is displayed based on cellular components (green), and molecular functions (blue).The number of proteins involved in each GO category were displayed on each terms. The color of each column represented the significance of each enriched categorize. D: The pathway enrichment analysis for these MTB-induced up-regulated proteins in MTB-infected BAM. The scatter plot visualized the pathway impact and enrichment results for all these significantly up-regulated proteins. Each point represented different metabolic pathways. The various color levels displayed different levels of significance of metabolic pathways from low (blue) to high (red). The size of each point represented the number of proteins involved in each corresponding metabolic pathway.

**Table 3.** Significantly enriched categories relevant to molecular function of eight MTB specifically activated proteins.

| GO ID | Description | Gene Number | pvalue | Genes |
| --- | --- | --- | --- | --- |
| GO:0008137 | NADH dehydrogenase (ubiquinone) activity | 2 | 0.000887 | P23935; P42028 |
| GO:0050136 | NADH dehydrogenase (quinone) activity | 2 | 0.000887 | P23935; P42028 |
| GO:0003954 | NADH dehydrogenase activity | 2 | 0.00098 | P23935; P42028 |
| GO:0016655 | oxidoreductase activity, acting on NAD(P)H | 2 | 0.001076 | P23935; P42028 |
| GO:0052642 | lysophosphatidic acid phosphatase activity | 1 | 0.002358 | A6H757 |
| GO:0016651 | oxidoreductase activity, acting on NAD(P)H | 2 | 0.003218 | P23935; P42028 |
| GO:0004427 | inorganic diphosphatase activity | 1 | 0.004711 | Q2KIV7 |
| **GO:0033721** | aldehyde dehydrogenase (NADP+)<br>activity | 1 | 0.004711 | E1BM93 |
| **GO:0052650** | NADP-retinol dehydrogenase<br>activity | 1 | 0.004711 | E1BM93 |
| **GO:0003993** | acid phosphatase activity | 1 | 0.007059 | A6H757 |
| **GO:0004033** | aldo-keto reductase (NADP)<br>activity | 1 | 0.009402 | E1BM93 |
| **GO:0008106** | alcohol dehydrogenase (NADP+)<br>activity | 1 | 0.009402 | E1BM93 |
| **GO:0016491** | oxidoreductase activity | 3 | 0.017809 | E1BM93; P23935;<br>P42028 |
| **GO:0003746** | translation elongation factor<br>activity | 1 | 0.030281 | P43896 |
| **GO:0051539** | 4 iron, 4 sulfur cluster binding | 1 | 0.037156 | P42028 |
| **GO:0016620** | oxidoreductase activity, acting on<br>NADP as acceptor | 1 | 0.043989 | E1BM93 |

“ $P < 0.05$ ”

The KEGG database was further analyzed to help determine the biological processes and functions of these eight MTB-activated proteins (Figure 4B). All these eight proteins were mainly related to energy metabolism represented by Oxidative phosphorylation and immune system represented by Complement and coagulation cascades (Figure 4B). Meanwhile, the significantly identified progress involved in response to pathogenic bacteria was Staphylococcus aureus infection (Figure 4B). In conclusion, under MTB infection challenges, dramatic activation of energy metabolisms was the main motion which was performed in BAM in response to MTB attacks.

### Functional analysis on up-regulated proteins which were specifically activated by MB infection

In the present study, to identify proteins only associated with MB-infection in BAM, protein expression profiles of three treatment (MTB-infection, MB-infection and normal BAM) were compared together. The hierarchical clustering of these up-regulated proteins showed that all of these significantly up-regulated proteins has similar trends that all these 51 up-regulated proteins were invoked to over-express only in MB-infected BAM (Figure 5C; Table 5).

**Figure 5.**
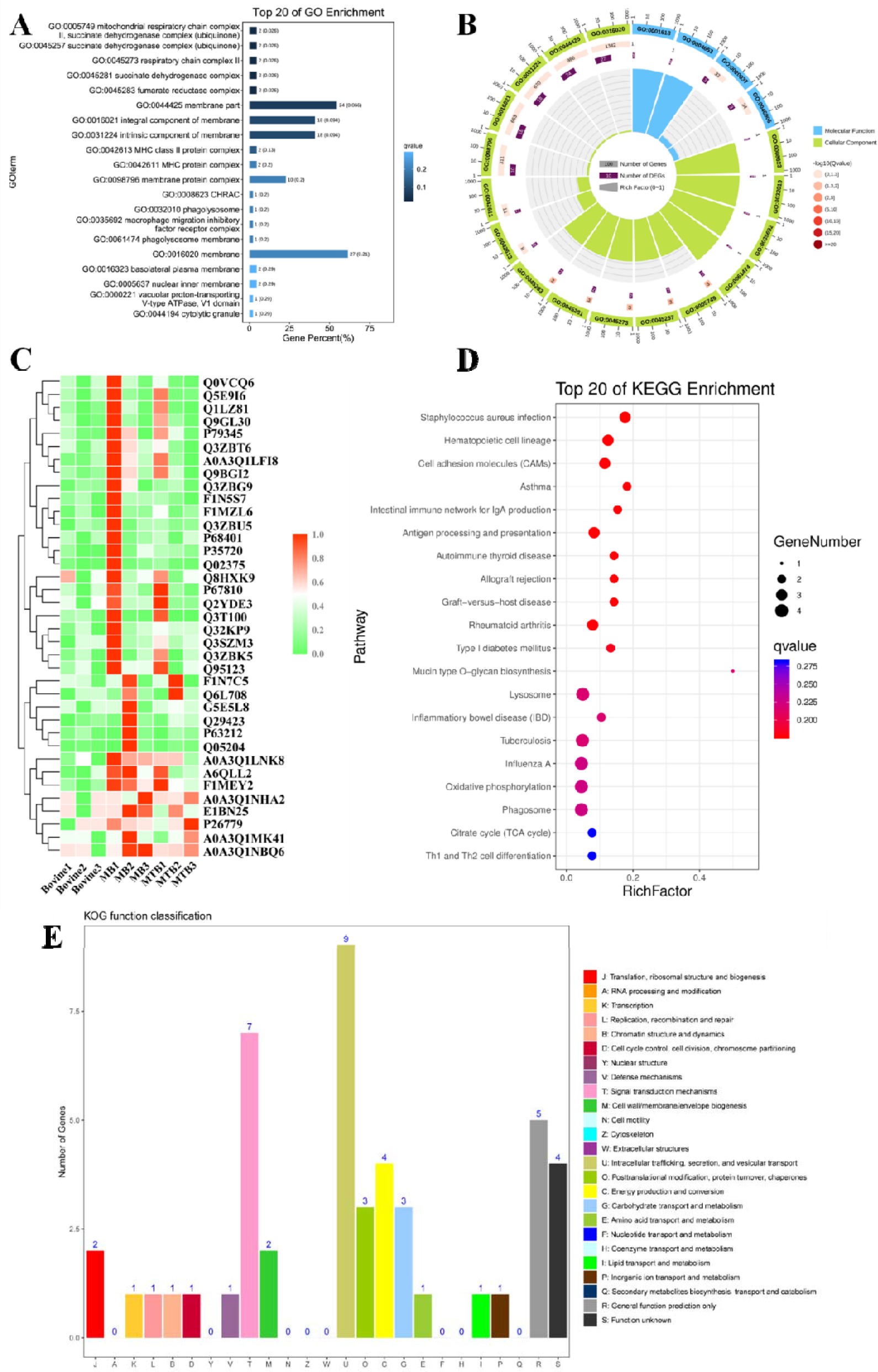
Functional annotation analysis on significanly up-regulated proteins associated with MB infection. A, B: GO enrichment analysis for 51 up-regulated proteins in BAM under MB infection challenges. The proteins were categorized according to the annotation of GO, and the number of each category is displayed based on cellular components (A), biological process and molecular functions (B).The number of proteins involved in each GO category were displayed on each terms. The color of each column represented the significance of each enriched categorize. C: Clustering analysis of proteins differentially expressed in these three treatment groups. Green indicated proteins with relative low expression abundance in the corresponding samples, whereas red indicated relative high expression abundance. D: The pathway enrichment results based on these MB-induced up-regulated proteins in BAM. The scatter plot visualized the pathway impact and enrichment results for all these 51 significantly up-regulated proteins. Each point represented different metabolic pathways. The various color levels displayed different levels of significance of metabolic pathways from low (blue) to high (red). The number of proteins involved in each pathway were shown by the size of each corresponding point. E: Functional classification of proteins by KOG analysis. The number of proteins in each KOG category was shown on the top on each column. J: Translation, ribosomal structure and biogenesis; A: RNA processing and modification; K: Transcription; L: Replication, recombination and repair; B: Chromatin structure and dynamics; D: Cell cycle control, cell division, chromosome partitioning; Y: Nuclear structure; V: Defense mechanisms; T: Signal transduction mechanisms; M: Cell wall/membrane/envelope biogenesis; N: Cell motility; Z: Cytoskeleton; W: Extracellular structures; U: Intracellular trafficking, secretion, and vesicular transport; O: Posttranslational modification, protein turnover, chaperones; C: Energy production and conversion; G: Carbohydrate transport and metabolism; E: Amino acid transport and metabolism; F: Nucleotide transport and metabolism; H: Coenzyme transport and metabolism; I: Lipid transport and metabolism; P: Inorganic ion transport and metabolism; Q: Secondary metabolites biosynthesis, transport and catabolism; R: General function prediction only; S: Function unknown

Subsequently, Gene Ontology enrichment analysis was carried out on all these significantly up-regulated proteins to elucidate the affected biological processes during the progress of pathogenic bacteria MB infection (Figure 5A, B; Table 4). For cellular component, a series of corresponding categorizes relevant to energy metabolisms, autophagy and inflammation were significantly identified, including mitochondrial respiratory chain complex II, succinate dehydrogenase complex (ubiquinone), respiratory chain complex II, phagolysosome, macrophage migration inhibitory factor receptor complex, phagolysosome membrane, autolysosome, NLRP3 inflammasome complex, AIM2 inflammasome complex and autophagosome (Figure 5A; Table 4). It suggested that MB infection could induce various changes in energy metabolisms, autophagy and inflammation of BAM. Additionally, three molecular functions relevant to oxidation-Reduction reactions were overrepresented among these 51 up-regulated proteins, including peroxidase activity, oxidoreductase activity and antioxidant activity (Figure 5B; Table 4). As imperative progresses, peroxidase activity, oxidoreductase activity and antioxidant activity perform important functions in BAM to defend pathogenic bacteria attacks, such as superoxide dismutase A (Shah et al., 2015). Remarkably, in biological progress categorizes, several terms associated with energy metabolisms, autophagy, defense response to bacteria and immune response were dramatically activated in BAM by MB infection, including defense response, hydrogen peroxide-mediated programmed cell death, immune response, mitochondrial electron transport, succinate to ubiquinone, negative regulation of inflammatory response, negative regulation of response to external stimulus, regulation of natural killer cell mediated immunity, respiratory burst after phagocytosis and response to wounding (Figure 5B; Table 4). All these GO enrichment analysis suggested that activation of energy metabolisms, autophagy, defense response to bacteria and immune response was the main response of BAM under pathogenic bacteria MB attacks.

**Table 4.**
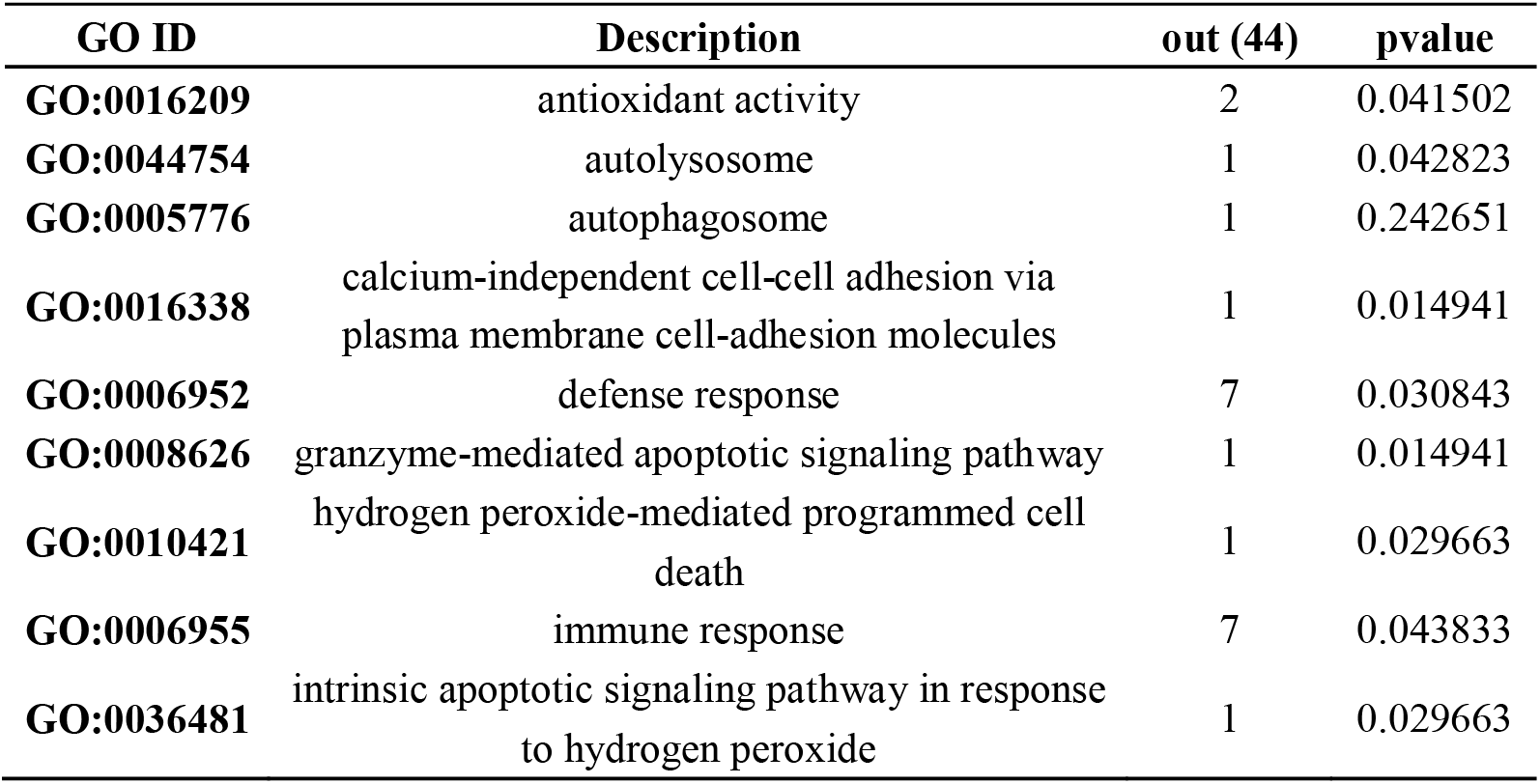
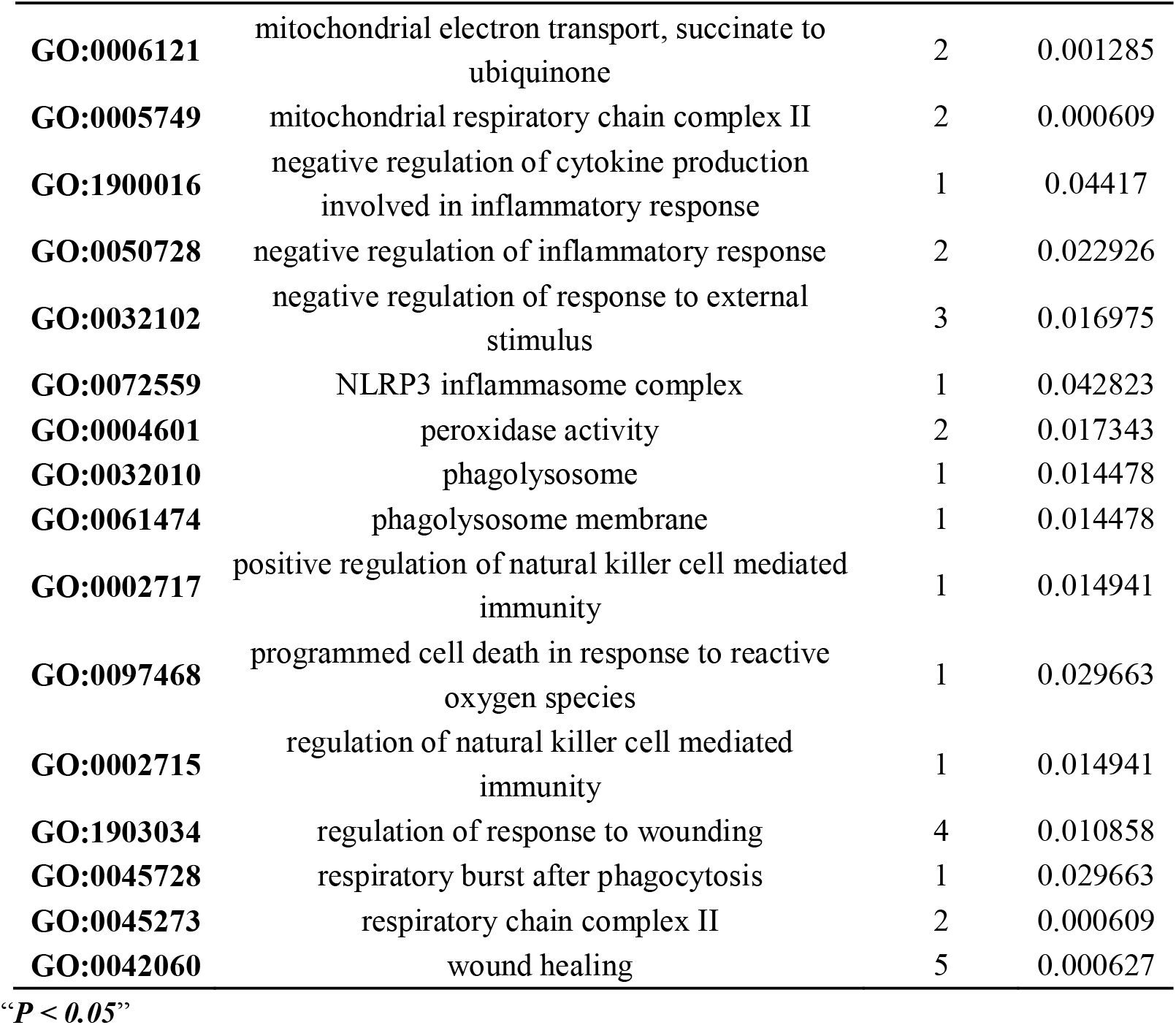
Significantly enriched Gene Ontology terms only in MB-infected bovine alveolar macrophages

**Table 5.** Key proteins activated by MB-infection.

| **Protein accession** | **Protein description** | **MB/Bovine Ratio** | **MB/Bovine P value** | **Subcellular localization** |
| --- | --- | --- | --- | --- |
| **A0A3S5ZPN6** | Scavenger receptor class B member 2 | 1.688 | 0.043886 | endoplasmic reticulum |
| **A5PJH7** | LOC788112 protein | 1.581 | 0.03298 | extracellular |
| **A6H6Y1** | BOLA-DQA1 protein | 1.519 | 0.036096 | peroxisome |
| **E1B726** | Plasminogen | 22.512 | 0.020516 | extracellular |
| **F1MGW6** | Uncharacterized protein | 2.404 | 0.00133542 | cytoplasm |
| **F1MNI5** | Prostaglandin G/H synthase 2 | 1.612 | 0.039212 | extracellular |
| **F1MX83** | Protein S100 | 2.48 | 0.039991 | cytoplasm |
| **F1MYR5** | Nitric oxide synthase | 1.615 | 0.015842 | cytoplasm,nucleus |
| **F1MZL6** | V-type proton ATPase subunit H | 1.705 | 0.037654 | cytoplasm |
| **F1N610** | Ig-like domain-containing protein | 1.607 | 0.031422 | extracellular |
| **G5E5L8** | Uncharacterized protein | 1.665 | 0.029085 | mitochondria |
| **P09428** | Interleukin-1 beta | 4.234 | 0.005355 | cytoplasm |
| **P26779** | Prosaposin | 2.484 | 0.023632 | extracellular |
| **P35720** | Succinate dehydrogenase cytochrome b560 subunit, mitochondrial | 1.89 | 0.043107 | plasma membrane |
| **P79345** | NPC intracellular cholesterol transporter 2 | 1.527 | 0.018958 | extracellular |
| **P81287** | Annexin A5 | 5.042 | 0.021295 | cytoplasm |
| **Q05204** | Lysosome-associated membrane glycoprotein 1 | 3.394 | 0.012726 | plasma membrane |
| **Q0VCQ6** | Programmed cell death 10 | 1.544 | 0.008831 | cytoplasm |
| **Q29423** | CD44 antigen | 1.62 | 0.018179 | plasma membrane |
| **Q3SZM3** | Cytochrome b-245 chaperone 1 | 1.552 | 0.034538 | mitochondria |
| **Q3T100** | Microsomal glutathione S-transferase 3 | 1.838 | 0.004157 | plasma membrane |
| **Q3ZBK5** | Tumor necrosis factor alpha-induced protein 8-like protein 2 | 2.805 | 0.013505 | cytoplasm |
| **Q6L708** | Claudin-1 | 1.564 | 0.011168 | plasma membrane |
| **Q8H XK9** | Apoptosis-associated speck-like protein containing a CARD | 1.531 | 0.007273 | cytoplasm |
| **Q95123** | Succinate dehydrogenase<br>[ubiquinone] cytochrome b<br>small subunit, mitochondrial | 1.736 | 0.028306 | mitochondria |
| **Q9BGI2** | Peroxiredoxin-4 | 1.619 | 0.044665 | extracellular |
| **F1MV85** | Polysaccharide biosynthesis<br>domain containing 1 | 0.664 | 0.025944 | cytoplasm |
| **P46168** | Beta-defensin 10 | 0.262 | 0.05 | extracellular |
| **A6QP29** | TBC1 domain family member<br>2A | 0.662 | 0.0034376 | cytoplasm |

“ $P < 0.05$ ”

As the database for protein orthologous classification, Cluster of Orthologous Groups of proteins analysis could contribute the understand of protein functions. We further analyzed these 51 up-regulated proteins with COG protein database to identify their specific function, the results showed that these proteins mainly involved in 17 of 25 KOG categories (Figure 5E). The highest frequency of occurrence of KOG terms are Signal transduction mechanisms, Intracellular trafficking, secretion, and vesicular transport and Energy production and conversion (Figure 5E). And the term Defense mechanisms was also involved among these MB-activated proteins (Figure 5E). Moreover, many identified proteins are involved in Posttranslational modification, protein turnover, chaperones and Carbohydrate transport and metabolism, indicating these functional classifications were also perform some functions in BAM in response to MB infection (Figure 5E).

Furthermore, according to KEGG pathway database analysis on these 51 proteins, the main biochemical metabolism and signal transduction pathways of BAM in response to MB infection has been described (Figure 5D). The results indicated that all annotated proteins were significantly mapped onto 18 KEGG pathways, especially Lysosome, Tuberculosis, Phagosome, Apoptosis, mTOR signaling pathway and Autophagy (Figure 5D). Remarkably, Tuberculosis which caused by MB infection was also significantly identified with Pvalue of 0.032 (Figure 5D). Additionally, biological progresses relevant to Autophagy and Apoptosis were still the major categorizes, which performed important function in defense response of BAM to pathogenic bacteria MB. And it has been proved that Lysosome and mTOR signaling pathway also perform important functions in defense of macrophage to MTB and MB infection (Bach et al., 2008; Wel et al., 2007; Yang et al., 2016; Yi et al., 2014; Fu et al., 2017).

Ultimately, we concluded that MB infection could cause changes in expression of proteins which involved in energy metabolism, autophagy, apoptosis, lysosome and inflammation. And the activation of these terms may be the major progresses of BAM to defend pathogenic MB attacks.

### MB-infection induced proteins were involved complex interaction network in bovine alveolar macrophages

Based on the analyses above, 29 proteins were identified as key proteins which were activated in BAM as being challenged by MB infection (Table 5). Among such 29 proteins, 26 proteins were activated by MB infection, whereas only 3 proteins F1MV85, P46168 and A6QP29 were suppressed by MB attacks in BAM (Table 5). Meanwhile, three proteins relevant to defense and autophagy were further verified by qRT-PCR. The results showed that all these three genes encoding these proteins were significantly induced to up-regulate in macrophage following MB infection, which further supported their important function in defending MB infection (Figure 6C). In the present study, we used STRING tool (version 10; Szklarczyk et al., 2011), which is directed to a database with known and predicted protein-protein interactions, to identify additional protein information for subsequent functional validation of these key MB infection-induced proteins (Figure 6A). Remarkably, the results depicted that plenty of protein-protein interactions occurred among these 29 key proteins (Figure 6A). However, for such condition, no proteins was interconnected with these down-regulated proteins (Figure 6A). On the contrary, in the further co-expression network, we observed a clearly negative correlation between those down-regulated proteins and up-regulated proteins, and forms a complex link between them (Figure 6B). And these proteins were mainly involved in various energy metabolisms-, autophagy- and immunity-related pathways, including Tuberculosis, Lysosome, Phagosome, Th17 cell differentiation and Oxidative phosphorylation (Table 6). We considered that there was a complex network of proteins, whose expression were altered in BAM as being challenged by MB infection to make defense responses.

**Figure 6.**
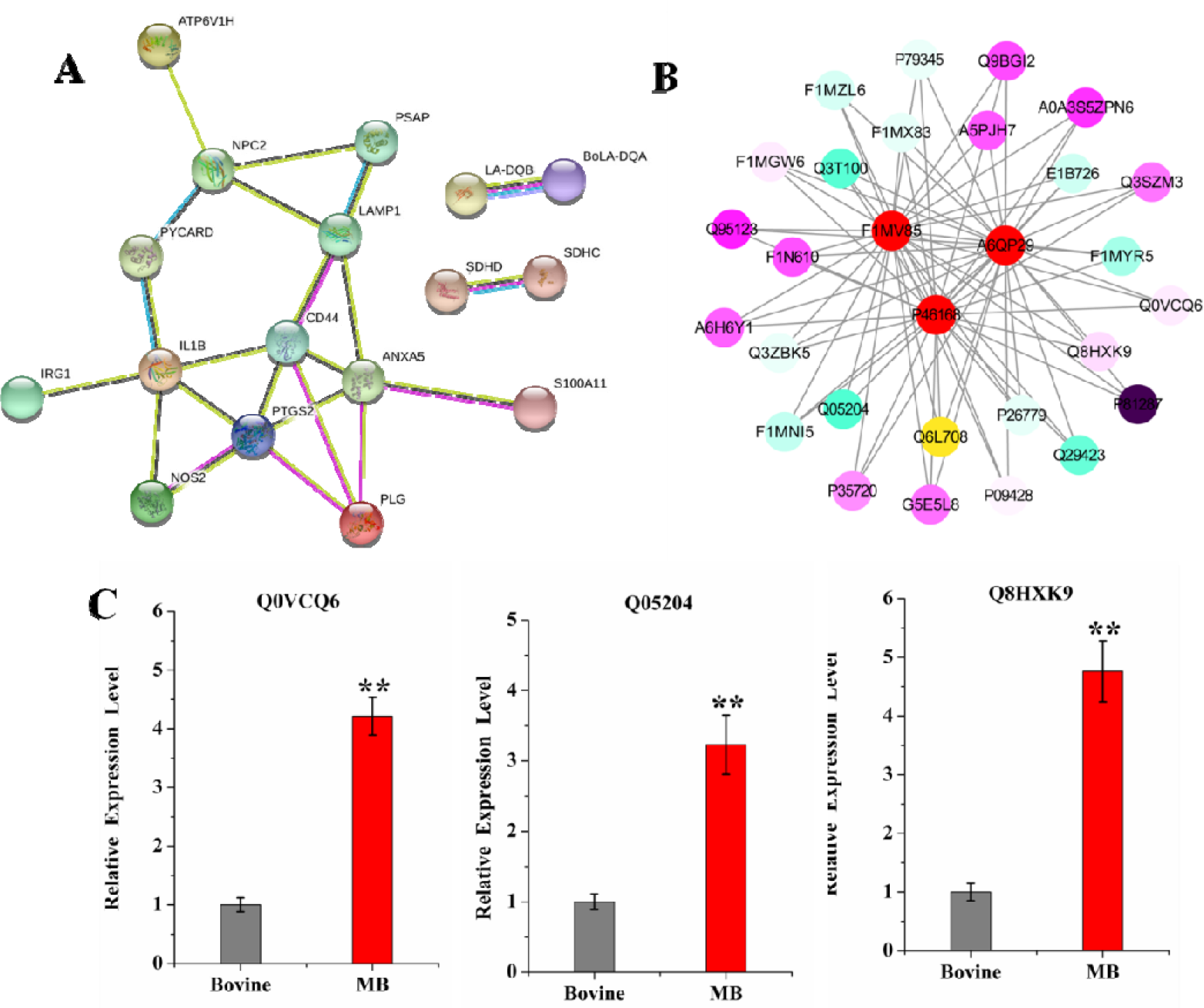
Complex interaction and co-expression network analysis on key MB-altered proteins. A: Global protein interaction network analysis of significantly MB-induced bovine proteins which involved in key pathways and GO terms. All the key differentially expressed proteins were submitted to the STRING tool (http://string.embl.de/) to predict protein-protein interaction network. B: Co-expression network analysis on significantly MB-induced bovine proteins which involved in key pathways and GO terms. The proteins shown in red circle represented three down-regulated proteins which was suppressed by MB infection in BAM. The various color levels displayed correlation level with its linked proteins from negative correlation (blue) to positive correlation (purple). C:qRT-PCR verification of important defense- and autophagy-related proteins induced by MB infection. The relative expression level were represented by the foldchange of 2^-□□Ct^ value.

**Table 6.** Key pathways of proteins activated by MB-infection.

| **#term**<br>**ID** | **term description** | **observed**<br>**gene count** | **false**<br>**discovery rate** |
| --- | --- | --- | --- |
| **bta05152** | Tuberculosis | 6 | 4.27E-06 |
| **bta04142** | Lysosome | 4 | 0.00022 |
| **bta04145** | Phagosome | 4 | 0.0005 |
| **bta04659** | Th17 cell differentiation | 3 | 0.0024 |
| **bta00190** | Oxidative phosphorylation | 3 | 0.0035 |
| **bta04672** | Intestinal immune network for IgA production | 2 | 0.0081 |
| **bta04657** | IL-17 signaling pathway | 2 | 0.0163 |
| **bta04064** | NF-kappa B signaling pathway | 2 | 0.0181 |
| **bta04668** | TNF signaling pathway | 2 | 0.0205 |

“ $P < 0.05$ ”

## Discussion

A recent study recognized that MTB and MB could invoke the responses of cattle and Homo sapiens, especially macrophages from two hosts (Daley et al., 2010). nutrition and a place for survival and reproduction for MTB and MB, and can also eliminate MTB and MB through apoptosis or autophagy (11-12). However, the different responses of Macrophages to both MTB and MB were still unclear. In this study, we used MTB- and MB-infected macrophage for in-depth proteomic analysis, and thus concluded that macrophages responded more strongly to MB than to MTB. During infection processes, M. tuberculosis could enhance its energy mechanism which may be better defense against MB and MTB infection. Meanwhile, macrophage could drive more active autophagy- and inflammatory-related progresses to defense MB infection, whereas MTB infection activates only 8 proteins involved in lysophosphatidic acid phosphatase activity, oxidoreductase activity, and ldehyde dehydrogenase (NADP+) activity. Furthermore, we found that various signaling pathways associated with autophagy- and inflammatory-related progresses were affected by MB infection which could contribute to the defense of macrophage to MB attacks. Collectively, our results provide novel insights into the different mechanisms of the pivotal MB- and MTB-macrophage interactions.

The interaction between M. bovis and macrophage results in chronic inflammatory and autophagy-related responses to prevent mycobacterial growth (Moraco et al., 2014). Autophagy is an important strategy for macrophages to kill MTB, and autophagic cell could not release intracellular MTB and MB components. As being ingested by phagocytosis, MTB will expose some antigen sites and release toxic substances. Stimulated macrophages can secrete cytokines and chemokines, thus initiating the occurrence of innate immunity (Wang et al. 2010). Macrophage can degrade the phagocytic MTB through acidic hydrolase by means of the fusion of phagocytic and lysosome, thus killing or inhibiting the growth of MTB in cells. Accordingly, MTB can inhibit phagocytic acidification, phagocytic and lysosomal fusion to avoid proteolytic enzyme hydrolysis and subsequent immune response events, which is the main strategy of MTB to avoid host cell clearance (Wel et al., 2007; Bach et al., 2008). Therefore, the expression of lysosomal related genes is also crucial for host pathogen elimination. In our study, we found that not only MTB infection can activate autophagy-associated proteins in macrophages, but MB infection can more strongly affect the expression of autophagy-associated proteins. In parallel, it was found that both MTB and MB infection could significantly invoke the expression of host lysosomal associated proteins, while MTB infection performed more significant influence on the activation of lysosomal associated proteins. Therefore, we proposed that MB could inhibit the expression of lysosomal and autophagy-related proteins such as F1MV85, P46168 and A6QP29, reduce the fusion of phagocytes and lysosomes, and thus weaken the effect on host autophagy, contributing to avoid its elimination by the immune response of host macrophages. Remarkably, researches have reported that lncRNA is involved in gene regulation of MTB infected hosts, and further affect autophagy related signaling pathways (Pawar et al., 2016). Therefore, we recognized that the change of lncRNA might be the main factor causing different host reactions when MTB and MB were infected, which needs further follow studies.

Finally, we found that these defense-related genes could response to MB infection through complex signaling networks, including NF-κB signaling pathway, IL-17 signaling pathway, Cytokine-cytokine receptor interaction, Inflammatory mediator regulation of TRP channels, Toll-like receptor signaling pathway and HIF-1 signaling pathway. It has been proved that TLR-2 pathway performed regulatory functions in immunomodulation through the export of agonist (Prados-Rosales et al., 2011; Athman et al., 2015). In parallel, recent researches have revealed that IL-17 is important for pro-inflammatory responses and chemokine regulation, and a stronger IL-17 response correlated with the severity of human diseases (Zuniga et al., 2012; Jurado et al., 2012). In vitro and in vivo generation of Th17 cells can lead to stronger IL-17 expression (Liang et al., 2006). Among these signaling, NF-κB and Toll-like receptor signaling pathway are strongly associated with activation of the host defense against MB infection, including various pro-inflammatory responses and release of anti-microbial effectors (Pilli et al., 2012; Liu et al., 2018; Jia et al., 2015). Researchers have demonstrated that MB is able to induce the production of TNF-α and IL in RAW264.7 cells and activates the NF-κB pathway through TLR2-mediated signaling transduction, which performed important roles in autophagy- and inflammatory-related progresses (Pilli et al., 2012; Liu et al., 2018). In contrast, the proteomic profiles of MTB-infected BAM did not exhibit the enrichment of these signaling pathways. It has been reported that MTB infection mainly suppresses MAPK signaling pathway and lipid metabolism pathway in macrophages, which is essential for regulating the expression of MTB-induced immunoregulatory molecules, such as TNF-α and IL (Shukla et al., 2017). Additionally, one study has demonstrated that MB can trigger autophagy- and inflammatory-related progresses mainly via NF-κB signaling pathway, IL-17 signaling pathway, Cytokine-cytokine receptor interaction, inflammatory mediator regulation of TRP channels, Toll-like receptor signaling pathways, which enhance our understanding of MB-mediated innate immune response of signaling pathway in MB-Macrophage interaction progress.

In summary, our research provides sufficient evidences that macrophage could make different responses to defend the MTB and MB infection, including various of autophagy and inflammatory-related progresses and signaling pathway. Furthermore, the results also indicated that MB could suppress the autophagy- and inflammatory-related proteins and signaling pathways to avoid host cell clearance for its colonization in host. Further study on the inhibitory effect of MB on host immune responses is essential for novel strategies to control such disease.

## Reference

Waters W R, Palmer M V. Mycobacterium bovivs infection of cattle and white-tailed deer: Translational research of relevance to human tuberculosis. Ilar J, 2015, 56(1): 26–43.

Minae A , Dong-Ryeol R , Jang W P , et al. ULK1 prevents cardiac dysfunction in obesity through autophagy-meditated regulation of lipid metabolism.[J]. Cardiovascular Research, 2017(10):1137.

Li X. Transmission of tuberculosis and drug resistant tuberculosis[D]. Fudan University , 2011.

Daley Charles, L . Update in tuberculosis 2009[J]. American Journal of Respiratory & Critical Care Medicine, 2010, 181(6):550.

Jing Ouyang, Jiayue Hu, Ji-Long Chen. lncRNAs regulate the innate immune response to viral infection[J]. Wiley Interdisciplinary Reviews: RNA, 2016.

Hmama Z , Pe?a-Díaz, Sandra, Joseph S , et al. Immunoevasion and immunosuppression of the macrophage by Mycobacterium tuberculosis[J]. Immunological Reviews, 2015, 264(1):220–232.

Moraco A H , Kornfeld H . Cell death and autophagy in tuberculosis[J]. Seminars in Immunology, 2014, 26(6):497–511.

Wel N V D , Hava D , Houben D , et al. M. tuberculosis and M. leprae translocate from the phagolysosome to the cytosol in myeloid cells.[J]. Cell, 2007, 129(7):1287-1298.

Horacio B ,Kadamba G. Papavinasasundaram , Dennis W , Zakaria H et al. Mycobacterium tuberculosis Virulence Is Mediated by PtpA Dephosphorylation of Human Vacuolar Protein Sorting 33B. Cell Host Microbe, 2008, 3(5): 316-322.

Fu Y , Xu X , Xue J , et al. Deregulated lncRNAs in B Cells from Patients with Active Tuberculosis[J]. Plos One, 2017, 12(1):e0170712.

Yi Z, Li J, Gao K, et al. Identification of differentially expressed long non-coding RNAs in CD4+ T cells response to latent tuberculosis infection. J Infect, 2014, 69(6): 558–568.

Yang X , Yang J , Wang J , et al. Microarray analysis of long noncoding RNA and mRNA expression profiles in human macrophages infected with Mycobacterium tuberculosis[J]. Rep, 2016, 6(1):38963.

Masters S L , Mielke L A , Cornish A L , et al. Regulation of interleukin-1β by interferon-γ is species specific, limited by suppressor of cytokine signalling 1 and influences interleukin-17 production[J]. EMBO reports, 2010.

Wynn T A , Chawla A , Pollard J W . Macrophage biology in development, homeostasis and disease[J]. Nature, 2013, 496(7446):445-455,c3.

Wang J X, Yang C. Mycobacterium tuberculosis and macrophages interaction research. Journal of Microbes and Infections, 2010, 5(3): 181–185.

Canaday D H , Wilkinson R J , Li Q , et al. CD4+ and CD8+ T Cells Kill Intracellular Mycobacterium tuberculosis by a Perforin and Fas/Fas Ligand-Independent Mechanism[J]. Journal of Immunology, 2001, 167(5):2734-42.

Sureka K , Sanyal S , Basu J , et al. Polyphosphate kinase 2: a modulator of nucleoside diphosphate kinase activity in mycobacteria[J]. Molecular Microbiology, 2010, 74(5):1187-1197.

Sánchez, Alejandro, Espinosa P , García, Teresa, et al. The 19?kDa *Mycobacterium tuberculosis* Lipoprotein (LpqH) Induces Macrophage Apoptosis through Extrinsic and Intrinsic Pathways: A Role for the Mitochondrial Apoptosis-Inducing Factor[J]. Clinical & Developmental Immunology, 2012, 2012:1–11. A. Casadevall, Evolution of intracellular pathogens, Ann. Rev. Micro 62 (2008) 19–33.

Schmittgen, T. D. & Livak, K. J. Analyzing real-time PCR data by the comparative C(T) method. Nat. Protoc. 3,1101–1108 (2008).

C.J. Cambier, K.K. Takaki, R.P. Larson, R.E. Hernandez, D.M. Tobin, K.B. Urdahl, L. Christine, Cosma, L. Ramakrishnan, Mycobacteria manipulate macrophage recruitment through coordinated use of membrane lipids, Nat 505 (2014) 218–222.

Pawar K , Hanisch C , Palma Vera S E , et al. Down regulated lncRNA MEG3 eliminates mycobacteria in macrophages via autophagy[J]. scientific Reports, 2016, 6:19416.

Bach H, Papavinasasundaram KG, Wong D, Hmama Z, Av-Gay Y. Mycobacterium tuberculosis virulence is mediated by PtpA dephosphorylation of human vacuolar protein sorting 33B. Cell Host Microbe. 2008 May 15;3(5):316-22. doi: 10.1016/j.chom.2008.03.008. PMID: 18474358.

Yi Z, Li J, Gao K, Fu Y. Identifcation of differentially expressed long non-coding RNAs in CD4+ T cells response to latent tuberculosis infection. J Infect. 2014 Dec;69(6):558-68. doi: 10.1016/j.jinf.2014.06.016. Epub 2014 Jun 27. PMID: 24975173; PMCID: PMC7112653.

Shah S , Cannon J R , Fenselau C , et al. A Duplicated ESAT-6 Region of ESX-5 Is Involved in Protein Export and Virulence of Mycobacteria[J]. Infection & Immunity, 2015:4349–4361.

Pilli, M.; Arko-Mensah, J.; Ponpuak, M.; Roberts, E.; Master, S.; Mandell, M.A.; Dupont, N.; Ornatowski, W.; Jiang, S.; Bradfute, S.B.; et al. TBK-1 promotes autophagy-mediated antimicrobial defense by controlling autophagosome maturation. Immunity 2012, 37, 223–234.

Liu, S.; Jia, H.; Hou, S.; Xin, T.; Guo, X.; Zhang, G.; Gao, X.; Li, M.; Zhu, W.; Zhu, H. Recombinant Mtb9.8 of Mycobacterium bovis stimulates TNF-α and IL-1 β secretion by RAW264.7 macrophages through activation of NF-κB pathway via TLR2. Sci. Rep. 2018, 8, 1928.

Jia, H.; Liu, S.; Wu, J.; Hou, S.; Xin, T.; Guo, X.; Yuan, W.; Gao, X.; Zhang, G.; Li, M.; et al. Recombinant TB9.8 of Mycobacterium bovis triggers the production of IL-12 p40 and IL-6 in RAW264.7 macrophages via activation of the p38, ERK, and NF-κB signaling pathways. Inflammation 2015, 38, 1337–1346.

Zuniga, J.; Torres-Garcia, D.; Santos-Mendoza, T.; Rodriguez-Reyna, T.S.; Granados, J.; Yunis, E.J. Cellular and humoral mechanisms involved in the control of tuberculosis. Clin. Dev. Immunol. 2012, 193923.

Jurado, J.O.; Pasquinelli, V.; Alvarez, I.B.; Pena, D.; Rovetta, A.I.; Tateosian, N.L.; Romeo, H.E.; Musella, R.M.; Palmero, D.; Chuluyan, H.E.; et al. IL-17 and ifn-gamma expression in lymphocytes from patients with active tuberculosis correlates with the severity of the disease. J. Leukoc. Biol. 2012 , 91, 991–1002.

Liang, S.C.; Tan, X.Y.; Luxenberg, D.P.; Karim, R.; Dunussi-Joannopoulos, K.; Collins, M.; Fouser, L.A. Interleukin (IL)-22 and IL-17 are coexpressed by Th17 cells and cooperatively enhance expression of antimicrobial peptides. J. Exp. Med. 2006, 203, 2271–2279.

Shukla S K , Shukla S , Chauhan A , et al. Pathway analysis of differentially expressed genes in Mycobacterium bovis challenged bovine macrophages[J]. Microbial Pathogenesis, 2017:343–352.

Prados-Rosales R, Baena A, Martinez LR, et al. Mycobacteria release active membrane vesicles that modulate immune responses in a TLR2-dependent VIRULENCE manner in mice. J Clin Invest. 2011;121(4):1471–1483.

Athman JJ, Wang Y, McDonald DJ, et al. Bacterial membrane vesicles mediate the release of Mycobacterium tuberculosis lipoglycans and lipoproteins from infected macrophages. J Immunol. 2015;195(3):1044–1053.

Moraco, Andrew H, Kornfeld, Hardy. Cell death and autophagy in tuberculosis[J]. Seminars in Immunology, 2014, 26(6):497-511.

